# Sex-specific brain structural predictors and outcomes of adolescent depression trajectories

**DOI:** 10.64898/2025.11.30.689896

**Authors:** Qingwen Ding, Divyangana Rakesh, Muskan Khetana, Sarah Whittle

**Author notes:** **Corresponding to:** Qingwen Ding, Centre for Youth Mental Health, The University of Melbourne, Parkville, Victoria, Australia.

## Abstract

**Objectives:** Depressive symptoms often emerge and increase during adolescence, yet the neural predictors and long-term brain outcomes associated with depressive trajectories remain poorly understood. This study adopts a trajectory-based approach to explore the temporal relationship between brain structure and dynamic changes in depressive symptoms across adolescence.

**Methods:** We used a large longitudinal adolescent neuroimaging sample across five annual waves (n = 11,862; mean age = 9.91 at baseline) from the ABCD study. First, growth mixture models were applied separately for males and females to identify distinct developmental trajectories of depressive symptoms. We then used linear mixed-effects models to identify which baseline brain structural features differentiated trajectory classes, and how these trajectories influenced subsequent brain structure.

**Results:** We identified high-decreasing, stable-low, and low-increasing trajectories in females, and relatively stable high, moderate, and low trajectories in males. No baseline brain regions were found to be associated with female trajectories, whereas smaller global brain measures (total subcortical volume, cortical volume, and surface area) predicted the high trajectory in males. Regarding brain outcomes, a low-increasing trajectory in females was associated with smaller global subcortical and cortical volumes, while high and moderate trajectories in males predicted thinner global cortices. Additional trajectory-related regional brain alterations were observed in both sexes.

**Conclusions:** These findings highlight distinct neurodevelopmental patterns linked to depressive symptom trajectories in males and females, underscoring the importance of a sex-specific and longitudinal approach to identifying neural risk and outcomes in adolescent depression.

## Introduction

Depressive symptoms often emerge and increase markedly during adolescence,^1^ a sensitive developmental period characterized by heightened brain plasticity.^2^ Atypical neurodevelopment during this time has been implicated in the onset and progression of adolescent depression.^3,4^ However, the predictive neural markers and long-term neural outcomes associated with adolescent depression symptoms remain poorly characterized. Such insights are crucial for improving early identification and informing more targeted intervention strategies.

According to the vulnerability hypothesis, atypical brain structure, induced by genetics and early adversity, may be a pre-existing risk factor for depression.^5,6^ Supporting this view, several prospective longitudinal studies have demonstrated that volumetric changes in limbic and striatal regions,^7,8^ as well as cortical thickness alterations in the frontal, temporal, insular, and occipital cortices during adolescence,^4,9,10^ are linked to an increased risk of depression. While these studies have made important contributions by highlighting the potential antecedent role of atypical brain structure in depression, they have consistently treated depression as a static trait. Indeed, depressive symptoms during adolescence are not stable, but instead evolve over time.^11^ Relying on cross-sectional symptom assessments risks misclassifying individuals and offers limited insight into the dynamic relationship between brain structure and depression development.

To address this limitation, only one study, to our knowledge, has examined brain structure in relation to changes in internalizing psychopathology during adolescence.^12^ This study found that lower cortical thickness in sensorimotor and temporal regions at ages 9–10 was associated with a steeper decline in future internalizing symptoms across adolescence. Although informative, it modeled symptom change at the population-averaged level and could not distinguish neural markers for individuals following different developmental patterns. Using person-centered approaches,^13^ extensive research has documented substantial variability in depressive symptom trajectories. Four distinct subgroups were commonly identified: stable-low, high-decreasing, low-increasing, and stable-high trajectories.^14,15^ These subgroups differ both in symptom severity and in long-term outcomes. For instance, adolescents following stable-high trajectory tend to experience the most persistent and severe symptoms and face greater risk for adverse developmental consequences compared to those with stable-low symptoms.^14^ Thus, the neural correlates underlying these trajectories are likely to differ, reflecting distinct pathways of neural vulnerability and resilience. Identifying trajectory-specific neurobiological predictors is essential for a more nuanced characterization of brain–behavior relationships and pinpointing early biological markers for risk trajectories.

Complementing the vulnerability hypothesis, the neurotoxicity hypothesis suggests that atypical brain structure could result from the cumulative effects of prolonged depressive symptoms over time, reflecting stress-related neurobiological alterations.^6,16^ A few studies have explored how depressive symptoms may longitudinally influence adolescent brain development within this framework.^17–20^ For instance, some studies reported that early depression onset was associated with subsequent reductions in cortical gray matter volume and cortical thinning,^17^ and depressive change (i.e., slope) was related to alterations in orbitofrontal cortex thickness.^18^ Another study, however, found no significant associations between depressive symptoms and later brain structure,^19^ highlighting inconsistencies in the literature. To date, only one study has investigated brain structural differences across individuals following distinct symptom trajectories. Schmaal et al.^20^ reported reduced surface area in the anterior cingulate and orbitofrontal cortex among adolescents with early-decreasing symptoms. However, this study was limited by a relatively small sample (n = 149) and focused on a narrow set of regions of interest (i.e., hippocampus, amygdala, and prefrontal cortices). Thus, larger, population-based samples are critically needed to identify how different trajectories may uniquely shape the adolescent brain.

Another critical gap concerns sex differences. Depressive symptom development diverges markedly by sex during adolescence, with boys showing relatively stable symptom levels while girls typically exhibit a gradual increase over time on an average level.^11,21^ Previous studies have also identified sex as a significant predictor of depression trajectories in the general population.^22–25^ Importantly, accumulating evidence indicates that both the neural predictors and neural outcomes of depression differ between females and males.^4,8,18,20^ As such, it is essential to investigate sex-specific developmental patterns and their associated neural pathways. To address this, large-scale studies are needed to explicitly model depressive trajectories and their neural correlates separately for males and females.

The present study utilized a large, representative longitudinal sample (n=11,862) from the Adolescent Brain Cognitive Development (ABCD) study. Our goal was to examine the brain structural predictors and outcomes of adolescent depression trajectories (Figure 1). First, we identified distinct developmental patterns of depressive symptoms using a person-centered growth mixture modeling (GMM) approach. Next, we applied a brain prediction model to identify baseline brain structural features that differentiate adolescents who later follow distinct depressive trajectories. We also implemented a brain outcome model to examine how depressive trajectories influence subsequent brain structure, while accounting for baseline neuroanatomy. Finally, to clarify sex-specific patterns in both depressive trajectories and their neural correlates, all analyses were conducted separately for female and male subgroups. These analyses were exploratory, and no a priori hypotheses were formulated.

**Figure 1.**
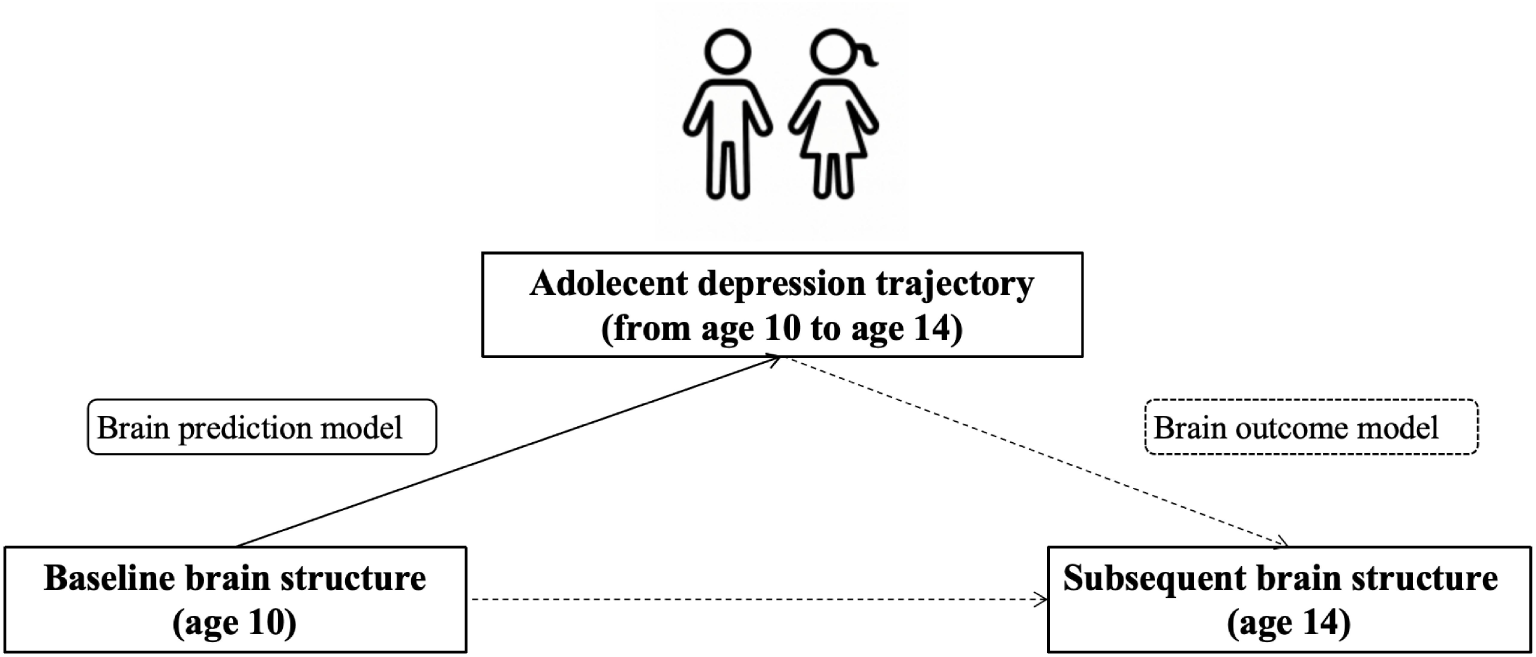
Research framework.

## Methods and materials

### Sample

The ABCD study is a longitudinal research project comprising a cohort of 11,862 participants, initially enrolled at ages 9 to 11 years, across 21 research sites in the United States. Participants with Child Behavior Checklist (CBCL) data at baseline were included in the analysis (n = 11,862, males = 6,185, females = 5,674). Data for the present study were obtained from the National Institute of Mental Health Data Archive (ABCD Release 5.0). Five waves of data were included: baseline, and follow-ups at one, two, three, and four years. Participants self-identified their race and ethnicity as Asian, Black, Hispanic, White, or Other. The study was conducted with appropriate ethics approval and informed consent.

### Depression symptoms

At each wave, depressive symptoms were assessed using the parent-reported DSM-oriented depression scale of the CBCL.^26^ This scale consists of 13 items, each rated on a 3-point scale: 0 (“Not true”), 1 (“Somewhat or sometimes true”), and 2 (“Very true or often true”). For the present analyses, we used T-scores, which are normed by sex and age, to better capture symptom changes from a subclinical or clinical perspective. T-scores of 65 or higher are considered in the clinical range, scores between 60 and 64 are classified as subclinical and may warrant monitoring, while scores below 60 fall within the typical range. The CBCL is one of the most widely used instruments for assessing emotional and behavioral problems in youth.^27^ Parent-reported depression scales have shown good validation in the ABCD study.^28^

### Brain structure

This study utilized structural magnetic resonance imaging (sMRI) data from both the baseline and four-year follow-up assessments. Neuroimaging data were acquired using three harmonized 3T MRI platforms (Siemens, Philips, and GE), employing standardized T1- and T2-weighted sequences. Preprocessing procedures involved distortion correction, intensity normalization, spatial alignment, and quality control. We first examined associations between symptom trajectories and four global brain structure measures: subcortical gray matter volume, total cortical volume, mean cortical thickness, and cortical surface area.

Subsequently, region-wise analyses were conducted to identify regional structural alterations. Specifically, we analyzed 16 subcortical gray volumes derived from FreeSurfer’s automated segmentation atlas, along with 68 cortical volumes, 68 cortical thickness measures, and 68 cortical surface areas based on the Desikan–Killiany atlas (both left and right hemisphere). Imaging acquisition and preprocessing followed standardized protocols.^29^ Cortical surface reconstruction and subcortical segmentation were performed by the ABCD Data Analysis and Informatics Resource Center (DAIRC) using FreeSurfer software (version 7.1.1). A complete list of brain regions included in the analysis can be found in Supplementary.

### Statistics

#### GMM

We first employed GMM, a person centered method, to identify distinct subgroups of individuals based on their developmental trajectories. GMM is a statistical technique that detects latent classes while accounting for individual variability within each class. GMM assigns participants to the most likely class using the posterior probabilities obtained. To aid model convergence, the intercept variance was freely estimated, while the slope parameter was fixed within classes. Missing data were handled using full-information maximum likelihood (FIML) estimation. To identify the optimal number of subgroups, models with one to five classes were fitted separately. Model fit was evaluated using the Akaike Information Criterion (AIC), Bayesian Information Criterion (BIC), sample-size adjusted BIC (SABIC), and entropy. Lower AIC, BIC, and SABIC values, combined with higher entropy, indicated better model fit and classification quality.^30^ A minimum class size of 5% of the total sample is recommended. All GMM analyses were conducted using the *lcmm* package in R (version 4.4.0).

#### Brain prediction model

Due to the difficulty of incorporating random effects in multinominal logistic regression, we employed linear mixed-effects models instead to test whether adolescents following different symptom trajectories differed in brain structure at baseline. In this model, standardized baseline brain measures, both global and regional indices, were used as dependent variables, with depression trajectory class as the primary independent variable. Family income, brain scanner, and parental education were included as fixed effects, while site and family were modeled as random effects. When modeling regional brain measures, the corresponding global measure was included as a fixed effect.

#### Brain outcome model

Linear mixed-effects models were also used to assess the influence of depression trajectory group on brain structure at the final assessment wave. Trajectory group was entered as the independent variable, with standardized brain measures at the final wave as the dependent variable. The corresponding baseline brain measure, family income, parental education, and scanner were included as fixed effects, and site and family were modeled as random effects. When modeling regional brain measures, the corresponding global measure was included as a fixed effect.

Linear mixed-effects models were fitted using the lme4 package, and p-values were obtained using the lmerTest package in R. All groups were compared with one another by changing the reference group.

#### Multiple comparison correction

False Discovery Rate (FDR) correction^31^ was applied across global brain measures as well as separately for each metric type: subcortical volume, cortical volume, cortical thickness, and cortical surface area for regional measures (pFDR < 0.05).

## Results

### Descriptive statistics

5,674 females and 6185 males were included in this study. Sample information and descriptive statistics for depression scores are indicated in Supplementary Table S1.

### Depression developmental trajectory across adolescence

The optimal model for the female group was the three-class solution, based on the lowest AIC, BIC, SABIC, and a high entropy (Supplementary Table S2). As shown in Figure 2, the three trajectories were labeled as high-decreasing (8.460%), stable-low (83.539%), and low-increasing (8.001%) symptoms. Females in the high-decreasing trajectory exhibited initially elevated symptom levels that declined over time from average subclinical to within the normal range. In contrast, females in the low-increasing trajectory showed a sharp increase in average symptoms, moving from initially normal levels to clinically significant levels across the follow-up period.

**Figure 2.**
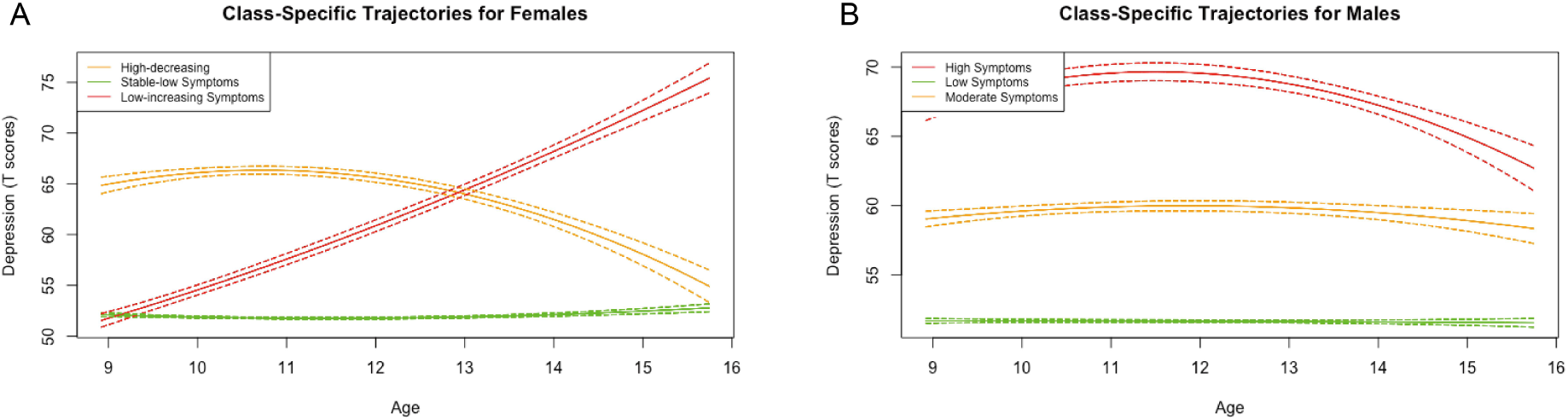
Depression trajectories for female and male adolescents.

For the male group, the three-trajectory model was also identified as the best fit (Supplementary Table S2). However, the trajectory patterns differed from those of the female group, consisting of high (5.078%), low (77.021%), and moderate (17.901%) symptom trajectories. Males in the high trajectory group exhibited elevated depressive symptoms in early adolescence, which gradually declined from the average subclinical range to below the subclinical threshold by mid-adolescence. Those in the moderate trajectory group showed average T-scores around 60 throughout the study period.

### Global brain structural predictors and outcomes for depressive trajectories

In females, no significant associations were found between global brain structures at baseline and depression trajectories (Table 1). In males, those following the high trajectory exhibited smaller total subcortical gray matter volume, cortical volume, and cortical surface area at baseline compared to those in the low trajectory group (*β* = –0.219 for subcortical volume; *β* = –0.193 for cortical volume; *β* = –0.193 for surface area) and moderate group (*β* = –0.231 for subcortical volume; *β* = –0.163 for cortical volume; *β* = –0.183 for surface area).

**Table 1.**
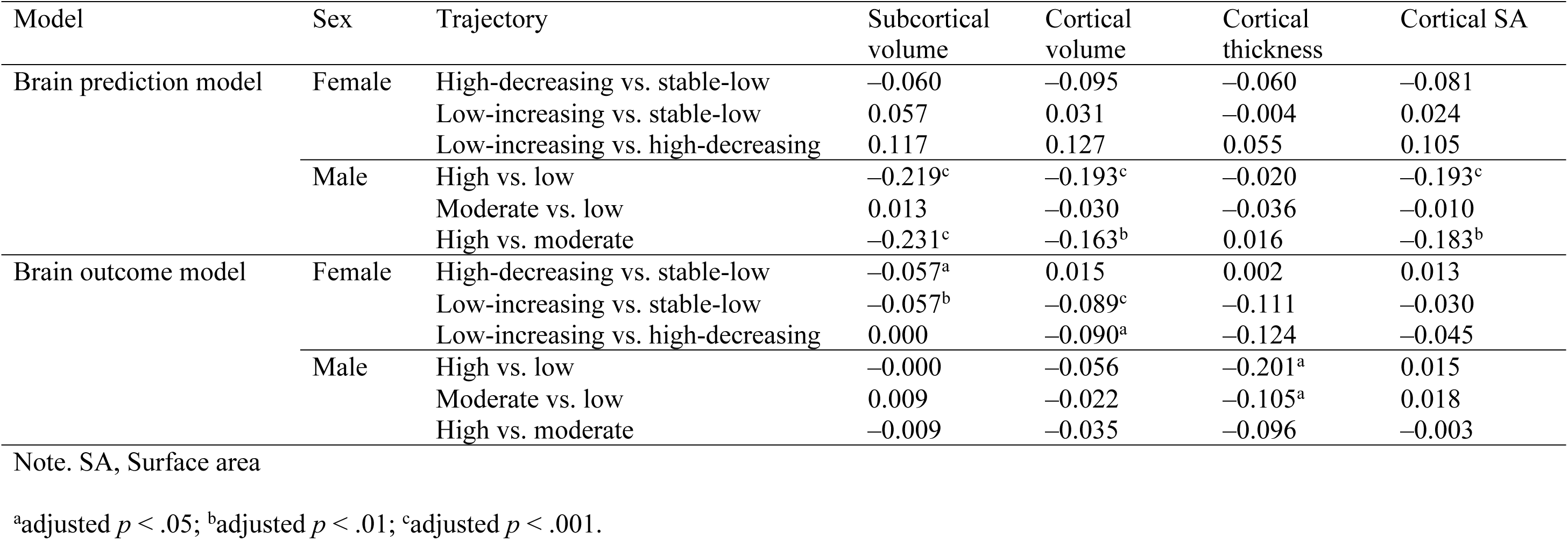
Global brain structure relations with depression trajectories.

| Model | Sex | Trajectory | Subcortical volume | Cortical volume | Cortical thickness | Cortical SA |
| --- | --- | --- | --- | --- | --- | --- |
| Brain prediction model | Female | High-decreasing vs. stable-low | −0.060 | −0.095 | −0.060 | −0.081 |
|  |  | Low-increasing vs. stable-low | 0.057 | 0.031 | −0.004 | 0.024 |
|  |  | Low-increasing vs. high-decreasing | 0.117 | 0.127 | 0.055 | 0.105 |
|  | Male | High vs. low | −0.219<sup>c</sup> | −0.193<sup>c</sup> | −0.020 | −0.193<sup>c</sup> |
|  |  | Moderate vs. low | 0.013 | −0.030 | −0.036 | −0.010 |
|  |  | High vs. moderate | −0.231<sup>c</sup> | −0.163<sup>b</sup> | 0.016 | −0.183<sup>b</sup> |
| Brain outcome model | Female | High-decreasing vs. stable-low | −0.057<sup>a</sup> | 0.015 | 0.002 | 0.013 |
|  |  | Low-increasing vs. stable-low | −0.057<sup>b</sup> | −0.089<sup>c</sup> | −0.111 | −0.030 |
|  |  | Low-increasing vs. high-decreasing | 0.000 | −0.090<sup>a</sup> | −0.124 | −0.045 |
|  | Male | High vs. low | −0.000 | −0.056 | −0.201<sup>a</sup> | 0.015 |
|  |  | Moderate vs. low | 0.009 | −0.022 | −0.105<sup>a</sup> | 0.018 |
|  |  | High vs. moderate | −0.009 | −0.035 | −0.096 | −0.003 |

Note. SA, Surface area
<sup>a</sup>adjusted $p < .05$ ; <sup>b</sup>adjusted $p < .01$ ; <sup>c</sup>adjusted $p < .001$ .

Regarding subsequent brain outcomes at the 4-year follow-up, females following low-increasing (*β* = –0.057) and high-decreasing (*β* = –0.057) trajectories had smaller total subcortical volumes compared to those following a stable-low trajectory. Additionally, those in the low-increasing trajectory exhibited smaller total cortical volume compared to both the high-decreasing (*β* = –0.090) and stable-low (*β* = –0.089) trajectory groups. Males following the high depression trajectory (*β* = –0.201) and moderate trajectory (*β* = –0.105) exhibited thinner mean cortical cortex at the four-year follow-up compared to low trajectory.

### Regional brain structural predictors and outcomes for depression trajectories

In females and males, we did not find differences in baseline regional brain structures between the three trajectories after controlling for global measures.

At the four-year follow-up, females in the high-decreasing group showed no significant differences in brain outcomes compared to the low-stable group. However, the low-increasing trajectory was associated with smaller volumes in the right hippocampus (*β* = –0.090) and 18 different cortical regions (*βs* = –0.058 to –0.130) compared to the low-stable trajectory. These cortical regions were broadly located in the inferior frontal gyrus (left pars opercularis and left pars triangularis), parietal lobe (left postcentral, right inferior parietal cortex, and left and right precuneus), temporal lobe (left superior temporal gyrus, right middle temporal cortex, right parahippocampal gyrus, and left and right fusiform gyrus), and occipital lobe (left and right pericalcarine cortex, left and right cuneus, left and right lingual gyrus, and right lateral occipital cortex). The low-increasing trajectory was also associated with smaller right lingual gyrus volume compared to the high-decreasing trajectory (*β* = –0.100). The estimates for those brain regions are shown in Figures 3A-3C.

**Figure 3.**
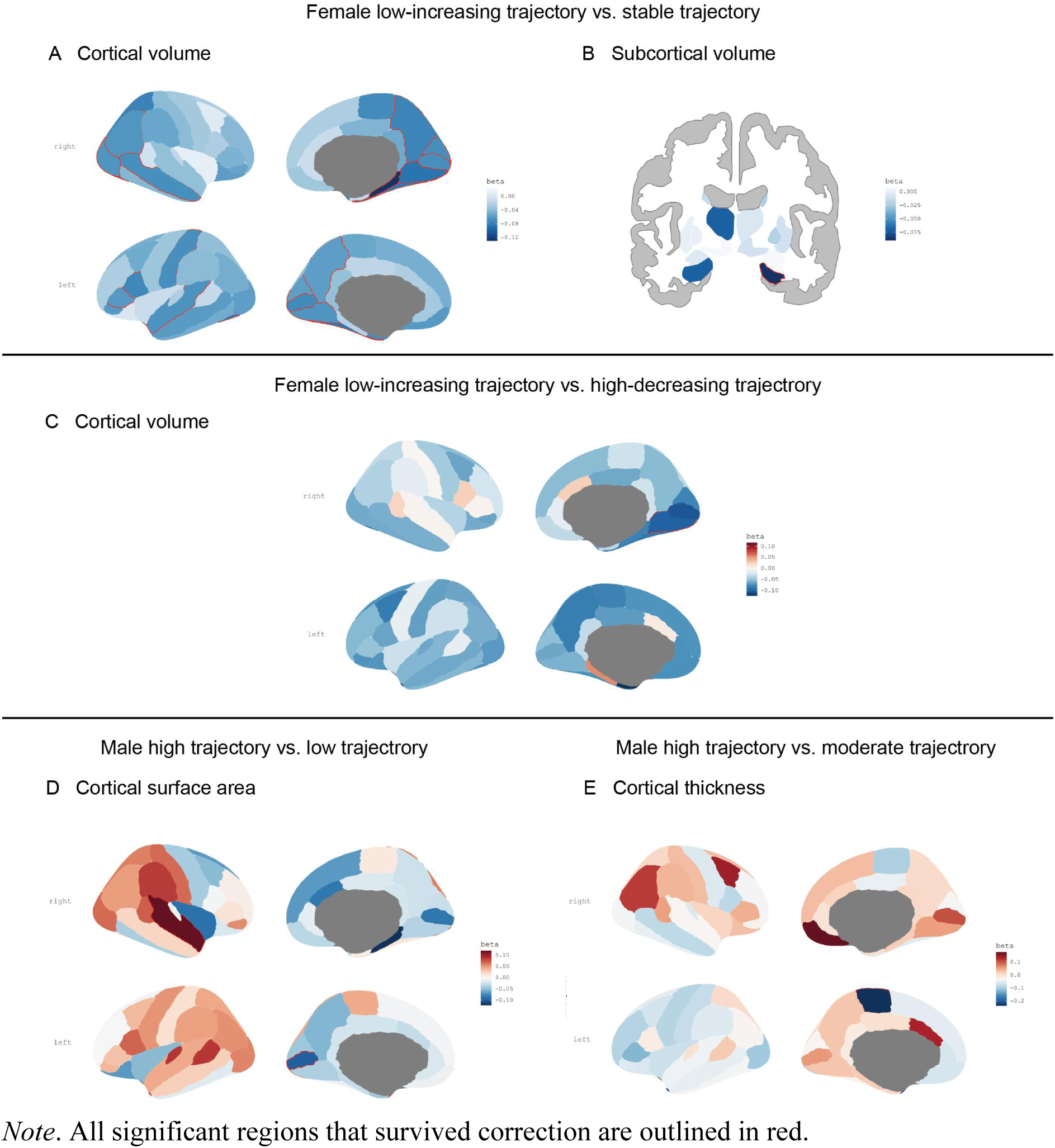
Regional brain structural outcomes for depression trajectories.

Compared to the low trajectory group, males in the high trajectory group showed a smaller surface area in the left pericalcarine cortex (*β* = –0.100) and a larger surface area in the right superior temporal cortex (*β* = 0.119) at the four-year follow-up. Males in the high trajectory also showed a significantly thinner cortex in the left-paracentral cortex than that in the moderate trajectory group (*β* = –0.239). The estimates for those brain regions are presented in Figures 3D and 3E.

## Discussion

This study provides insights into whether brain structural differences in early adolescence can predict future depression trajectories and whether these trajectories influence later brain outcomes, with consideration of sex differences. We found a high-risk trajectory in males that was characterized by persistently high symptoms during adolescence, while in females, two potentially high-risk trajectories were observed, marked by either increasing or decreasing symptom patterns. Smaller global brain volume and surface area predicted the high symptom trajectory in males, but not in females. In terms of brain outcomes, an increasing trajectory was prospectively associated with smaller global subcortical and cortical volumes in females, whereas in males, persistent high and moderate symptoms were associated with subsequent thinner global cortices. Beyond global metrics, some trajectory-related regional brain differences were also identified in both sexes. These findings enhance our understanding of the dynamic interplay between brain development and depression during adolescence and underscore the importance of sex-specific approaches in identification and prevention strategies.

Our study extended previous literature by using a large adolescent sample to explicitly model developmental trajectories separately for males and females,^22–25^ revealing distinct patterns across adolescence. For males, three distinct, non-intersecting trajectories characterized by high, moderate, and low symptom levels were identified. This pattern aligns with previous research suggesting a relatively stable pattern of average depressive symptoms in boys throughout adolescence.^11,21^ The emergence of high-symptom trajectory in our study is consistent with previous research showing that a subset of adolescent males had a tendency to experience a chronic form of depression.^23^ In contrast, the trajectories among females showed greater fluctuation, with patterns identified as high-decreasing, low-increasing, and stable-low. Our findings support some previous observations that females are more likely to follow trajectories marked by increasing depressive symptoms.^14,25^ Possible explanations for this sex difference may include significant cyclical hormonal fluctuations,^32^ and increased rates of social adversities (e.g., bullying),^33^ potentially contributing to the emergence of new or worsening depressive symptoms in some adolescent girls.

Males with smaller subcortical and cortical volumes, as well as reduced cortical surface area in early adolescence, were more likely to follow a high-symptom trajectory compared to those in the low or moderate trajectory. This pattern of globally reduced volume and surface area across widespread brain regions aligns with previous findings linking adolescent depression to impairments in multiple cognitive and affective domains.^34,35^ It also supports earlier prospective research showing associations between reduced global brain volume and surface area and general psychopathology in youth.^12^ However, the present study extends this work by examining these associations within the context of dynamic depressive symptom trajectories over time. The results suggest that global brain volume and surface area may be predictive markers to help distinguish males in the high-risk trajectory class (high symptoms) from those in low-risk trajectory classes (moderate and low symptoms). In contrast, we did not identify biological markers that predicted depressive risk trajectories in females. The observed sex difference may stem from divergent developmental patterns of depression. According to our results, males following high trajectories tend to exhibit early and stable symptom patterns, making structural brain differences easier to detect. In contrast, high-risk females tend to follow more dynamic symptom trajectories, which may obscure clear associations with brain structure.

Consistent with the neurotoxicity hypothesis, we found evidence that prolonged depression symptoms may alter brain structure in both males and females. This may be driven by chronic activation of the hypothalamic-pituitary-adrenal axis and elevated levels of pro-inflammatory cytokines, both well-documented in the pathophysiology of depression^36^, which can impair neuronal integrity and inhibit neurogenesis.^6,37^ Unlike previous studies of brain structure in adults and children,^38–40^ which typically report region-specific changes or solely subcortical changes, our results revealed more widespread alterations—reflected in reduced subcortical volume, cortical volume, and cortical thickness at a global level. This is likely because adolescence is a sensitive period for brain maturation, characterized by processes such as synaptic pruning and myelination.^41^ Depression, which is associated with chronic dysregulation of the stress response system,^42^ during this critical window may disrupt these processes, potentially leading to premature or excessive pruning across multiple regions, which may manifest as globally reduced brain volumes and thickness. The neuroanatomical outcomes longitudinally influenced by risk trajectories differed by sex: in females, alterations were primarily observed in subcortical and/or cortical volume, whereas in males, changes were more prominent in cortical thickness. Such sex-specific effects are likely shaped by the interplay of hormonal influences and divergent brain maturation patterns that differ markedly between sexes during adolescence. For example, estradiol levels in girls have been associated with reductions in gray matter volume,^43^ and girls typically show earlier declines in brain volume compared to boys.^44^ However, the underlying mechanisms are likely complex, and current evidence remains limited, warranting further investigation.

Neuroanatomical outcomes may vary based on the duration and severity of depressive symptoms associated with different trajectories. In females, the emerging risk group (i.e., the low-increasing trajectory) was associated with more widespread structural alterations, affecting both cortical and subcortical volumes. In contrast, the high-decreasing group, despite elevated symptom levels in early adolescence, exhibited structural effects limited to subcortical volume. Notably, the low-increasing group had smaller total cortical volume than both the high-decreasing and stable-low groups, suggesting a unique vulnerability associated with the emergence of new symptoms during adolescence. Among males, both the high and moderate symptom trajectories were linked to significant future lower thickness, with no significant differences observed between the two groups. This finding suggests both risk trajectories lead to global cortical thinning, and that even moderate but persistent depressive symptoms may contribute to structural brain alterations in males. These sex-specific findings may help explain why both low-increasing trajectories^45^ and moderate and high symptom classes^46^ have been found associated with the poorest psychosocial outcomes in previous research.

After controlling for global brain measures, significant associations between depression trajectories and future regional brain alterations were observed. These findings suggest that depression is linked not only to widespread structural changes but also to focal, region-specific effects, consistent with previous research on internalizing psychopathology.^12^ In females, those following a low-increasing trajectory showed reduced volume in widespread regions involved in language, motor function, sensory integration, memory, and visual processing, areas that have also been implicated in adolescent major depressive disorder.^47^ Many of these regions are higher-order association areas that mature later in adolescence,^48^ and may therefore be particularly susceptible to the effects of emerging depressive symptoms. Notably, females in the low-increasing group exhibited reduced volume in the right lingual gyrus, a region involved in emotional visual processing, compared to both the stable low and high-decreasing groups, suggesting this region may represent a unique structural outcome associated with the low-increasing trajectory. This may indicate that the lingual gyrus is especially sensitive to newly emerging depressive symptoms during adolescence. In contrast, regional structural alterations were more restricted in males, and primarily involved regions related to sensory and motor processing, including the left pericalcarine cortex, right superior temporal cortex, and left paracentral cortex.

This study offers several notable strengths. First, we innovatively applied a person-centered approach to capture heterogeneity in the association between brain structure and distinct depressive developmental trajectories. Second, we examined not only brain predictors of symptom trajectory but also trajectory-related brain outcomes, shedding light on the temporal precedence of the brain-depression relationship. Third, the large sample size enabled us to model males and females separately, providing a nuanced insight into sex-specific pathways. Nonetheless, a few limitations should be acknowledged. First, because our focus was on neural differences between depressive symptom trajectories, we were unable to account for prior depressive symptoms in a way that would support strong causal inference. Future research could benefit from combining cross-lagged models with trajectory models to more rigorously examine causal relationships. Second, our findings are based on an adolescent sample; thus, additional studies involving children and adults are needed to determine whether these patterns generalize across developmental stages. Finally, we relied solely on parent-reported data in this study. Incorporating multiple informants in future research would provide a more comprehensive understanding of depressive trajectories and their neural correlates.

## Conclusion

This study examined predictive biomarkers that distinguish depressive symptom trajectories and assessed brain outcomes associated with these trajectories in adolescent males and females. Specifically, male depression trajectories appear to be more predictable using global brain structural measures compared to females. Both sexes exhibited significant brain alterations associated with depressive risk trajectories, with particularly notable changes observed in the low-increasing trajectory among females and the high trajectory among males. These findings contribute to a deeper understanding of the neural mechanisms underlying depression development and may help inform sex-specific prevention and intervention strategies.

