## Supplementary 1 for "Sex-specific brain structural predictors and outcomes of adolescent depression trajectories"

Supplementary Material

**Regions of interest**

*Subcortical regions*  Thalamus proper (L), Caudate (L), Putamen (L), Pallidum (L), Hippocampus (L), Amygdala (L), Nuclear accumbens area (L), Ventral diencephalon (L), Thalamus proper (R), Caudate (R), Putamen (R), Pallidum (R), Hippocampus (R), Amygdala (R), Nuclear accumbens area (R), Ventral dien cephalon (R)

*Cortical regions*  Caudal middle frontal (L), frontal pole (L), lateral orbitofrontal cortex (L), medial orbitofrontal cortex (L), paracentral lobule (L), pars opercularis (L), pars orbitalis (L), pars triangularis (L), precentral gyrus (L), rostral middle frontal gyrus (L), superior frontal gyrus (L), inferior parietal lobule (L), postcentral gyrus (L), precuneus cortex (L), superior parietal lobule (L), supramarginal gyrus (L), banks of superior temporal sulcus (L), entorhinal cortex (L), fusiform gyrus (L), inferior temporal gyrus (L), middle temporal gyrus (L), parahippocampal gyrus (L), superior temporal gyrus (L), temporal pole (L), transverse temporal cortex (L), cuneus cortex (L), lateral occipital cortex (L), lingual gyrus (L), pericalcarine cortex (L), caudal anterior cingulate (L), isthmus cingulate cortex (L), posterior cingulate cortex (L), rostral anterior cingulate cortex (L), insular (L), caudal middle frontal (R), frontal pole (R), lateral orbitofrontal cortex (R), medial orbitofrontal cortex (R), paracentral lobule (R), pars opercularis (R), pars orbitalis (R), pars triangularis (R), precentral gyrus (R), rostral middle frontal gyrus (R), superior frontal gyrus (R), inferior parietal lobule (R), postcentral gyrus (R), precuneus cortex (R), superior parietal lobule (R), supramarginal gyrus (R), banks of the superior temporal sulcus (R), entorhinal cortex (R), fusiform gyrus (R), inferior temporal gyrus (R), middle temporal gyrus (R), parahippocampal gyrus (R), superior temporal gyrus (R), temporal pole (R), transverse temporal cortex (R), cuneus cortex (R), lateral occipital cortex (R), lingual gyrus (R), pericalcarine cortex (R), caudal anterior cingulate (R), isthmus cingulate cortex (R), posterior cingulate cortex (R), rostral anterior cingulate cortex (R), insular (R).

Table S1. Sample size and descriptive statistics.

| Time point | Sample size | Mean age (SD) | Mean depression raw score (SD) |
| --- | --- | --- | --- |
| Baseline | 11862 | 9.91(0.63) | 1.27(2.01) |
| One-year follow up | 11203 | 10.90(0.64) | 1.39(2.19) |
| Two-year follow up | 10899 | 12.00(0.67) | 1.47(2.24) |
| Three-year follow up | 10099 | 12.90(0.65) | 1.68(2.47) |
| four-year follow up | 4679 | 14.10(0.68) | 1.76(2.70) |

Table S2 Fit indices for growth mixture modeling.

| Sex |  | Model fit | | | | | Class proportions | | | | |
| --- | --- | --- | --- | --- | --- | --- | --- | --- | --- | --- | --- |
| Male | k | Log likelihood | AIC | BIC | SABIC | entropy | class1 | class2 | class3 | class4 | class5 |
|  | 1 | -68425.61 | 136861.2 | 136894.9 | 136879.0 | 1.00 | 100.00 |  |  |  |  |
|  | 2 | -67212.52 | 134443.0 | 134503.6 | 134475.0 | 0.91 | 86.64 | 13.36 |  |  |  |
|  | **3** | **-66684.53** | **133395.1** | **133482.6** | **133441.2** | **0.90** | **5.08** | **77.02** | **17.90** |  |  |
|  | 4 | -66684.53 | 133403.1 | 133517.5 | 133463.5 | 0.58 | 0 | 18.43 | 76.49 | 5.08 |  |
|  | 5 | -66685.28 | 133413.8 | 133555.1 | 133488.4 | 0.62 | 0 | 75.78 | 5.11 | 19.11 | 0 |
| Female | 1 | -62858.30 | 125726.6 | 125759.8 | 125743.9 | 1.00 | 100.00 |  |  |  |  |
|  | 2 | -61521.75 | 123061.5 | 123121.3 | 123092.7 | 0.93 | 12.02 | 87.98 |  |  |  |
|  | **3** | **-60779.66** | **121585.3** | **121671.7** | **121630.4** | **0.87** | **8.46** | **83.54** | **8.00** |  |  |
|  | 4 | -60779.67 | 121593.3 | 121706.3 | 121652.3 | 0.50 | 9.01 | 8.64 | 82.36 | 0 |  |
|  | 5 | -60779.66 | 121601.3 | 121740.8 | 121674.1 | 0.86 | 8.04 | 0 | 8.46 | 83.50 | 0 |
